# Ultra-high field MRI reveals mood-related circuit disturbances in depression: A comparison between 3-Tesla and 7-Tesla

**DOI:** 10.1101/459479

**Authors:** Laurel S Morris, Prantik Kundu, Sara Costi, Abigail Collins, Molly Schneider, Gaurav Verma, Priti Balchandani, James W Murrough

## Abstract

Ultra-high field 7-Tesla (7T) MRI has the potential to advance our understanding of neuropsychiatric disorders, including major depressive disorder (MDD). To date, few studies have quantified the advantage of resting state functional MRI (fMRI) at 7T compared to 3-Tesla (3T). We conducted a series of experiments that demonstrate the improvement in temporal signal-to-noise ratio (TSNR) of a multi-echo fMRI protocol with ultra-high field 7T, compared to 3T MRI in healthy controls (HC). We also directly tested the enhancement in ultra-high field 7T fMRI signal power by examining the ventral tegmental area (VTA), a small midbrain structure that is critical to the expected neuropathology of MDD but difficult to discern with standard 3T MRI. We demonstrate 200-300% improvement in TSNR and resting state functional connectivity coefficients provided by ultra-high field 7T fMRI compared to 3T, indicating enhanced power for detection of functional neural architecture. A multi-echo based acquisition protocol and signal denoising pipeline afforded greater gain in signal power at ultra-high field compared to classic acquisition and denoising pipelines. Furthermore, ultra-high field fMRI revealed mood-related neuro-circuit disturbances in patients with MDD compared to HC, which were not detectable with 3T fMRI. Ultra-high field 7T fMRI may provide an effective tool for studying functional neural architecture relevant to MDD and other neuropsychiatric disorders.

## Introduction

In the past 30 years, functional magnetic resonance imaging (MRI) has provided unprecedented insight into the neural mechanisms of neuropsychiatric disorders in humans, including major depressive disorder (MDD). Work with 3-Tesla (3T) fMRI has revealed the functional architecture of key neural systems that contribute to the neuropsychopathology of MDD, including aberrant connectivity of limbic and reward-related networks that subserve mood regulation(1–4). The next generation of ultra-high field 7-Tesla (7T) MRI magnet has just been approved by the US Food and Drug Administration (FDA) for clinical use and while it has significant potential to advance our understanding of neuropsychiatric disorders, there remain concerns about data quality and systematic comparisons between 7T and 3T functional MRI remain scarce (5–8).

Ultra-high field 7T MRI benefits from increased signal to noise ratio (SNR)(5, 7), enhanced amplitude and percent of signal change(5, 8, 9) and increased susceptibility induced and blood oxygen level dependent (BOLD) contrast, all important for functional and spectroscopic applications(6). However, higher field strengths also produce more B0 inhomogeneities and susceptibility artifacts, which lead to geometric distortion, signal drop-out and signal pile-up. These distortions are most severe around tissue-air boundaries and more subtle susceptibility distortions can arise from paranasal sinuses and bone(10). Ventral portions of the brain, including subcortical and midbrain structures important for mood regulation, are particularly affected by susceptibility distortions and signal drop-out due to their proximity to bone and sinuses(11, 12). These limitations have led to concerns regarding the utility of ultra-high field MRI for examining ventral regions relevant to neuropsychiatric disorders. Shorter echo-time (TE), thinner slices and parallel imaging can combat some of these issues by reducing intra-voxel inhomogeneity and through-plane dephasing(6, 9, 13). We have developed and implemented an ultra-high field 7T functional MRI scanning protocol with multiple TE’s, thin slices, multi-band acquisition and TE-dependent physiological denoising methods to improve signal power detection and temporal SNR (TSNR) for *in vivo* imaging.

The application of ultra-high field 7T functional MRI to the study of neuropsychiatric disorders such as MDD remains scarce. MDD is one of the world’s largest public health issues to date, affecting approximately 300 million people and representing the leading cause of disability worldwide(14). The current lack of widely effective preventative interventions demonstrates in part our limited understanding of the etiology and biological mechanisms of this complex disorder. One of the core symptoms of MDD is anhedonia, a markedly diminished response to pleasure, which is characterized by underlying dysfunction of limbic and reward-related neural systems centered on VTA, nucleus accumbens and anterior cingulate cortex (ACC)(15–17). VTA hyperactivity and hyperconnectivity has been demonstrated in a well-validated preclinical model of depression that is characterized by the onset of anhedonic behaviors (18–20). VTA connectivity, however, has not been fully explored in humans with MDD, partly due to limited feasibility of discerning VTA at current standard 3T field strength (11, 12).

Here, we detail a series of experiments that demonstrate the functional MRI signal quality of a multi-echo scanning protocol at ultra-high field 7T MRI, and compare it to 3T MRI. Our first aim was to compute the gain in TSNR with ultra-high field 7T MRI, both across the whole brain and in particular neural regions relevant to MDD and related neuropsychiatric disorders. We subsequently examined functional connectivity of three essential fronto-striatal-midbrain circuits that mediate cognitive, limbic and motor functions(21–23) using ultra-high field 7T as compared to 3T MRI. Our second aim was to characterize the expected improvement in signal power more directly in healthy controls (HC) and patients with MDD by examining the functional connectivity of the VTA. We specifically selected the VTA due to its critical role in the expected neuropathology of MDD coupled with the difficulty in its delineation with standard 3T MRI. We hypothesized that whole-brain TSNR and circuit-specific functional connectivity would be improved at ultra-high field 7T compared to 3T, and further that 7T functional MRI would reveal VTA circuit differences between MDD and HC.

## Methods

### Participants

All subjects were recruited at the Mood and Anxiety Disorder Program (MAP) at Icahn School of Medicine at Mount Sinai. All eligible participants between the ages of 18-65 underwent the Structured Clinical Interview for DSM-V Axis Disorders (SCID-V) by a trained rater to determine any current or lifetime psychiatric disorder(24). Subjects were excluded if they had an unstable medical illness, history of neurological disease, history of schizophrenia or other psychotic disorder, neurodevelopmental/neurocognitive disorder, substance use disorder (SUD) within the past 2 years, any contraindications to MRI, or positive urine toxicology on the day of scan. HC subjects were free from any current or lifetime psychiatric disorder. All participants were free of antidepressant medication or other psychotropic medication for at least 4 weeks (8 weeks for fluoxetine) prior to data collection. Inclusion criteria for MDD subjects included having MDD as their primary presenting problem and being in a current major depressive episode (MDE). In all subjects, depressive symptom severity was measured with the Montgomery–Åsberg Depression Rating Scale (MADRS)(25) and severity of anhedonia was measured with the Snaith-Hamilton Pleasure Scale (SHAPS)(26). Subjects underwent either 3T or 7T MRI and a subset of individuals completed both 3T and 7T. All data was collected under Institutional Review Board (IRB)–approved written informed consent. All subjects were compensated for their time.

### MRI acquisition and preprocessing

Functional MRI data was collected with 3T Siemens Magnetom Skyra and 7T Siemens Magnetom scanners using a 10-minute multi-echo multi-band echo-planar imaging (EPI) pulse sequence with the following parameters at 3T: 3mm isotropic resolution, 45 slices, TR/TE’s= 882 / 11,29.7, 48.4, 67.1ms, MB=5, iPAT acceleration factor=2, 664 frames, flip=45, field of view= 560×560, pixel bandwidth=2085; and at 7T: 2.5mm isotropic resolution, 50 slices, TR/TE’s= 1850 / 8.5, 23.17, 37.84, 52.51, iPAT acceleration factor=3, 300 frames, flip=70, field of view= 640×640, pixel bandwidth=1786. Anatomical data were collected at 3T and at 7T with a twice magnetization-prepared rapid gradient echo (MP2RAGE) sequence for improved T1-weighted contrast and spatial resolution (27, 28), with the following parameters at 3T: 1mm isotropic resolution, 56 slices, TR/TE= 4000/1.9ms, field of view= 184 x 160, bandwidth=250; and at 7T: 0.7mm isotropic resolution, 60 slices, TR/TE= 6000/3.62ms, field of view= 240 x 320, bandwidth= 300.

Functional data were processed and denoised for motion and physiological artefacts using freely available multi-echo independent components analysis (ME-ICA, https://bitbucket.org/prantikk/me-ica)(29). This is explained in detail elsewhere (29, 30). Briefly, the ME-ICA package exploits the property that BOLD percent signal change is linearly dependent on TE, a consequence of T2* decay (29, 30). ME-ICA decomposes multi-echo functional MRI data into independent components, and computes the TE dependence of the BOLD signal for each component, as measured using the pseudo-*F*-statistic, Kappa, with components that scale strongly with TE having high Kappa scores(29). Non-BOLD components are identified by TE independence measured by the pseudo-*F*-statistic, Rho. As such, components are categorized as BOLD or non-BOLD based on their weightings measured by Kappa and Rho values, respectively. By removing non-BOLD components, data are denoised for motion, physiological and scanner artefacts in a robust manner based on physical principles(29, 30). Denoised functional data were coregistered with T1 and normalized to a standard template with Advance Normalization Tools (ANTs, http://stnava.github.io/ANTs/) software using diffeomorphic symmetric normalization transformation (SyN) registration(31, 32).

Data from TE2 (29.7ms at 3T and 23.17 at 7T) were processed using a standard denoising pipeline implemented in AFNI software (afni_proc.py(33)), which included whole brain voxel timeseries despiking, correction for slice timing offsets, image realignment to a middle slice for volume-to-volume rigid body correction, masking, regression of demeaned motion parameters plus derivatives and censorship of time points with motion >0.2mm, and spatial smoothing with a full width half maximum kernel of 6mm (these procedures including smoothing were performed for the single-echo data only). While this does not represent an optimal single-echo functional MRI protocol, it provides a useful illustrative comparison.

### Data analysis

Demographic data and clinical characteristics were summarized and compared using independent t-tests, including for the clinician-administrated scale of depression severity (MADRS) and the self-report measure of anhedonia (SHAPS) (Table 1).

**Table 1.** Demographic and Clinical Characteristics. Abbreviations: Number of subjects (N), 3-Tesla (3T), 7-Tesla (7T), Major Depressive Disorder (MDD), healthy control (HC), Montgomery–Åsberg Depression Rating Scale (MADRS), Snaith-Hamilton Pleasure Scale (SHAPS), standard deviation (SD).

|  | 3T |  | 7T |  |
| --- | --- | --- | --- | --- |
|  | MDD | HC | MDD | HC |
| N | 15 | 17 | 10 | 17 |
| Age at enrollment, years (mean ± SD) | 40.7±11.560 | 37.4±10.137 | 34.3 ± 11.186 | 38.41 ±11.9 |
| Male (frequency, percent) | 9, 60% | 12, 70.6% | 6, 60% | 9, 52.9% |
| White (frequency, percent) | 8, 53.3% | 8, 47.1% | 6, 60% | 6, 46.2% |
| Hispanic ethnicity (frequency, percent) | 6, 40% | 1, 5.9% | 2, 20% | 2, 15.4% |
| College degree, at least 2-yr (frequency, percent) | 12, 80% | 15, 88.3% | 8, 80% | 10, 76.9% |
| Employed, at least part-time (frequency, percent) | 10, 66.7% | 14, 82.4% | 7, 70% | 11, 84.6% |
| Married (frequency, percent) | 1, 6.7% | 3, 17.6% | 0% | 2, 15.4% |
| Age at first episode (mean ± SD) | 23.6±13.695 | - | 22 ± 12.055 | - |
| Years since first episode (mean ± SD) | 17.1±10.575 | - | 12.3 ± 6.913 | - |
| Number of episodes (median, range) | 1; 39 | - | 2.5; 48 | - |
| Duration of current MDE, months (median, range) | 120; 498 | - | 13.5; 190 | - |
| Recurrent MDD (frequency, percent) | 4, 26.7% | - | 6, 60% | - |
| SHAPS Score (mean ± SD) | 35.27 ± 7.206 | 17.1 ± 4.293 | 39.6 ± 5.966 | 18.7 ± 4.9 |
| MADRS score (mean ± SD) | 28.4 ± 4.687 | 0.824 ± 1.131 | 29.8 ± 7.569 | 0.5 ± 0.9 |

For both 3T and 7T functional MRI data, temporal SNR (TSNR) was computed as the mean/standard deviation of the voxel signal timecourses of the raw EPI data for each TE and of the denoised data before coregistration and normalization across the whole brain. Using the functional MRI data normalized to template space, TSNR was also computed for the 7 regions that subserve cognitive and behavioural processes relevant to psychiatric research: ventral tegmental area, nucleus accumbens, amygdala, subgenual ACC, dorsal ACC, ventromedial prefrontal cortex (PFC), and dorsolateral PFC. The anatomical boundaries and definitions of these regions are described in detail elsewhere(34).

With the normalised functional MRI data at 3T and 7T, a 7×7 cross-correlation matrix was computed for the 7 regions of interest using standard Pearson correlation of the mean timecourse of each region, followed by Fisher’s R-to-Z transformation of the correlation coefficient (35). Percent differences were computed between 3T and 7T Fisher Z-transformed correlation coefficients.

In addition, seed-to-voxel functional connectivity maps were computed for the VTA, ventromedial PFC, dorsolateral PFC and SMA to examine networks relevant to depression as well as three cortical-basal ganglia circuits relevant to broader psychiatric research (23, 34). For seed-to-voxel functional connectivity maps, whole-brain statistical cluster-correction was performed with AFNI’s 3dClustSim, correcting for the instance of false positives due to multiple comparisons and spatial autocorrelation (36). For experiment 1, the correction threshold for the 3T data was calculated at voxelwise p<0.001 and Cluster>27 voxels for alpha<0.05. At 7T it was at voxelwise p<0.001 and Cluster>919 voxels for alpha<0.05.

For the examination of patients compared to controls, we examined seed-to-seed functional connectivity, computed as the Pearson correlation between averaged timecourses for VTA and nucleus accumbens; VTA and subgenual ACC; and VTA and dorsal ACC. These were entered into independent samples t-test to compare between groups at both 3T and 7T. For this p< 0.017 was considered significant, with bonferonni correction for 3 comparisons. Seed-to-seed functional connectivity for these regions was additionally correlated with duration of current episode and the depressive symptom of anhedonia. Finally, on an exploratory basis, VTA seed-to-whole-brain functional connectivity maps were compared at both 3T and 7T using independent samples t-test. We examined the data using a more lenient threshold: for the 3T data was calculated at voxelwise p<0.01 and Cluster>104 for alpha<0.05; and at 7T, voxelwise p<0.01 and Cluster>4645 for alpha<0.05.

**Figure 1.**
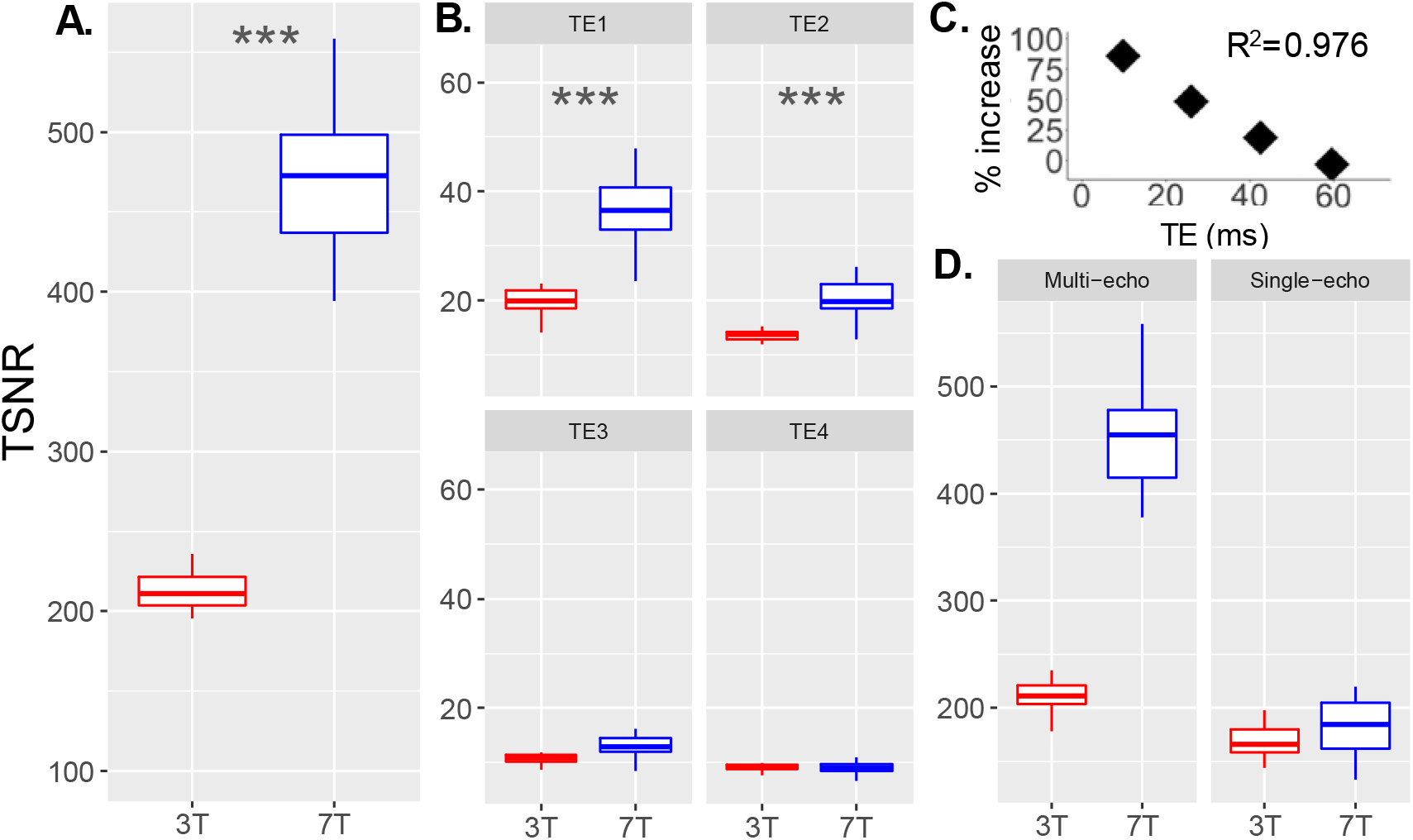
Higher temporal signal to noise ratio (TSNR) with ultra-high field 7-Tesla (7T) functional magnetic resonance imaging (MRI) scales with echo time (TE) and is boosted by multi-echo based denoising. A. TSNR plotted for whole brain at 3-Tesla (3T) and 7T. B. TSNR plotted for 7T at 3T for 4 different TE’s (TE1=shortest, TE4=longest). C. Percent increase in TSNR at 7T compared to 3T scales with TE (Linear fit, adjusted R^2^=0.976, p=0.008). D. TSNR plotted for whole brain after multi-echo acquisition and denoising and traditional single-echo acquisition and denoising. ***p<=0.001

## Results

### Experiment 1: Signal power at high field MRI in healthy controls

Seventeen healthy control (HC) participants were recruited and scanned on 3T (5 female, age= 37.4 ±10.1) and 17 on 7T (8 female, age= 38.41 ±11.9). Six of the 17 subjects completed both 3T and 7T scans (1 female, age=36.8 ±11.6). See Table 1 for clinical and demographic details.

#### Shorter and multiple TE’s contribute to improved SNR at high field

Whole brain multi-echo denoised EPI data had 227.98% higher TSNR at 7T compared to 3T (t_(32)_=11.6, p=2.7×10^−12^, Figure 1A). Raw EPI TSNR was also examined for each TE. Whole brain TSNR was increased at 7T compared to 3T at shorter TEs but not longer TEs (TE1= 176.29%, t_(32)_=5.5, p=3.6×10^−5^; TE2= 140.96%, t_(32)_=4.0, p=0.001; TE3=112.87%, t_(32)_=1.8, p=0.096; TE4= 92.32%, t_(32)_=-1.6, p=0.144; Figure 1B). The percent improvement in TSNR from 3T to 7T was linearly related to TE (adjusted R^2^=0.98, p=0.008, Figure 1C).

Multi-echo EPI data acquisition and denoising methods improved TSNR compared to conventional single echo acquisition and denoising (afni_proc.py with TE2) by 23.0% at 3T (t_(16)_=2.8, p=0.012) and by 148.4% at 7T (t_(16)_=15.7, p=3.8×10^−11^; Figure 1D), indicating increased SNR gain produced by ME-ICA at 7T compared to 3T. Motion was not significantly different between 3T and 7T (millimeter displacement, p=0.37; RPY degrees, p=0.68).

The same pattern of improved TSNR at 7T compared to 3T for whole brain denoised EPI and individual TE EPI data was observed when comparing 5 MDD subjects who completed 3T and 7T (whole-brain TSNR at 3T= 211.6 ±25.1; 7T= 425.3 ±42.5; p=0.006) and 5 HC subjects who completed both 3T and 7T (whole-brain TSNR at 3T= 200.1 ±52.6; 7T= 507.8 ±110.2; p=0.005; Supp Fig 1).

#### Higher field strength yields improved SNR in ventral regions relevant to psychiatric research

Regional TSNR was examined for 7 brain regions relevant to psychiatric research. Similar to whole brain improvement in TSNR at 7T compared to 3T, all 7 regions showed increased TSNR at higher field, including ventral structures close to sinuses and bone (Fig 2A, Supp Table 1, Supp Fig 3). The same pattern of improved TSNR at 7T compared to 3T for regional data was observed when comparing 5 HC subjects who completed both 3T and 7T (Supp Fig 1).

**Figure 2.**
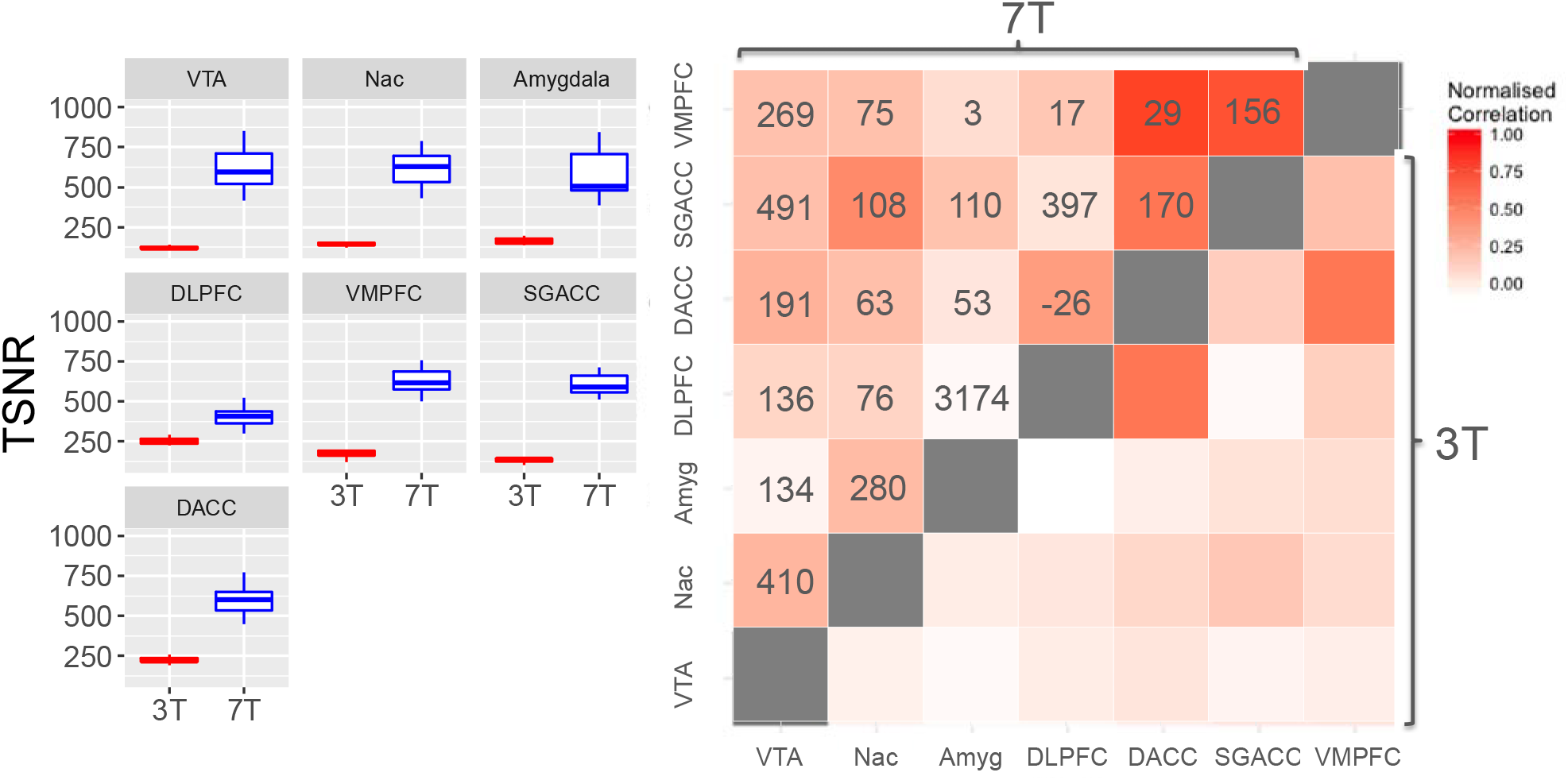
Improvement in ultra-high field 7-Tesla (7T) functional magnetic resonance imaging (MRI) signal power and cross-correlation coefficients throughout the brain. A. Temporal signal to noise ratio (TSNR) plotted for regions of interest at 3-Tesla (3T) and 7T. B. Matrix of Fisher Z-transformed cross-correlation coefficients between the same brain regions at 3T (bottom right) and 7T (top left). Values indicate percent improvement in correlation coefficient at 7T compared to 3T. Abbreviations: Ventral tegmental area (VTA), nucleus accumbens (Nac), amygdala (Amyg), dorsolateral prefrontal cortex (DLPFC), ventromedial prefrontal cortex (VMPFC), subgenual anterior cingulate cortex (SGACC), dorsal anterior cingulate cortex (DACC).

Correlation coefficients between these 7 brain regions were compared between 3T and 7T. Normalised coefficients were higher at 7T compared to 3T (Figure 2B). The pattern of connectivity was similar at 3T and 7T. For example, the subgenual cingulate cortex had the highest correlation coefficient (functional connectivity) with NAc and lowest with dlpfc at both 3T and 7T (Figure 2B). The average percent improvement from 3T to 7T across this diverse cross-correlation matrix was 300.9%. The same pattern was observed in the same HC and MDD subjects scanned on both 3T and 7T (Supp Table 2). To determine whether 3T and 7T correlation patterns were statistically similar, we examined normalized cross-correlation coefficients at 3T and 7T for 470 approximately same sized regions based on the Harvard-Oxford Atlas (37), finding significant correlation between 3T and 7T 470×470 cross-correlation matrices (R=0.38, p<0.0001).

Other cortical – basal ganglia circuits relevant to psychiatric research were compared between 3T and 7T. Three broadly dissociable putative circuits were examined, thought to subserve discrete motor, cognitive and limbic functions, based on SMA, dlpfc and vmpfc connectivity, respectively(23, 34). At 3T, cortical – basal ganglia circuits were distinguishable, reflecting dissociable motor (SMA, motor cortex, putamen), cognitive (dlpfc, MD thalamus) and limbic (vmpfc, nucleus accumbens, some VTA) subcircuits (Fig 3). However, at 7T the precision of identification of each subcircuit was heightened (Fig 3, Table 2 for statistics). For example, motor circuit nodes SN, posterior STN, ventrolateral thalamus were apparent; cognitive circuit node dorsal caudate; and limbic circuit nodes of VTA, locus coeruleus were apparent (Fig 3).

**Figure 3.**
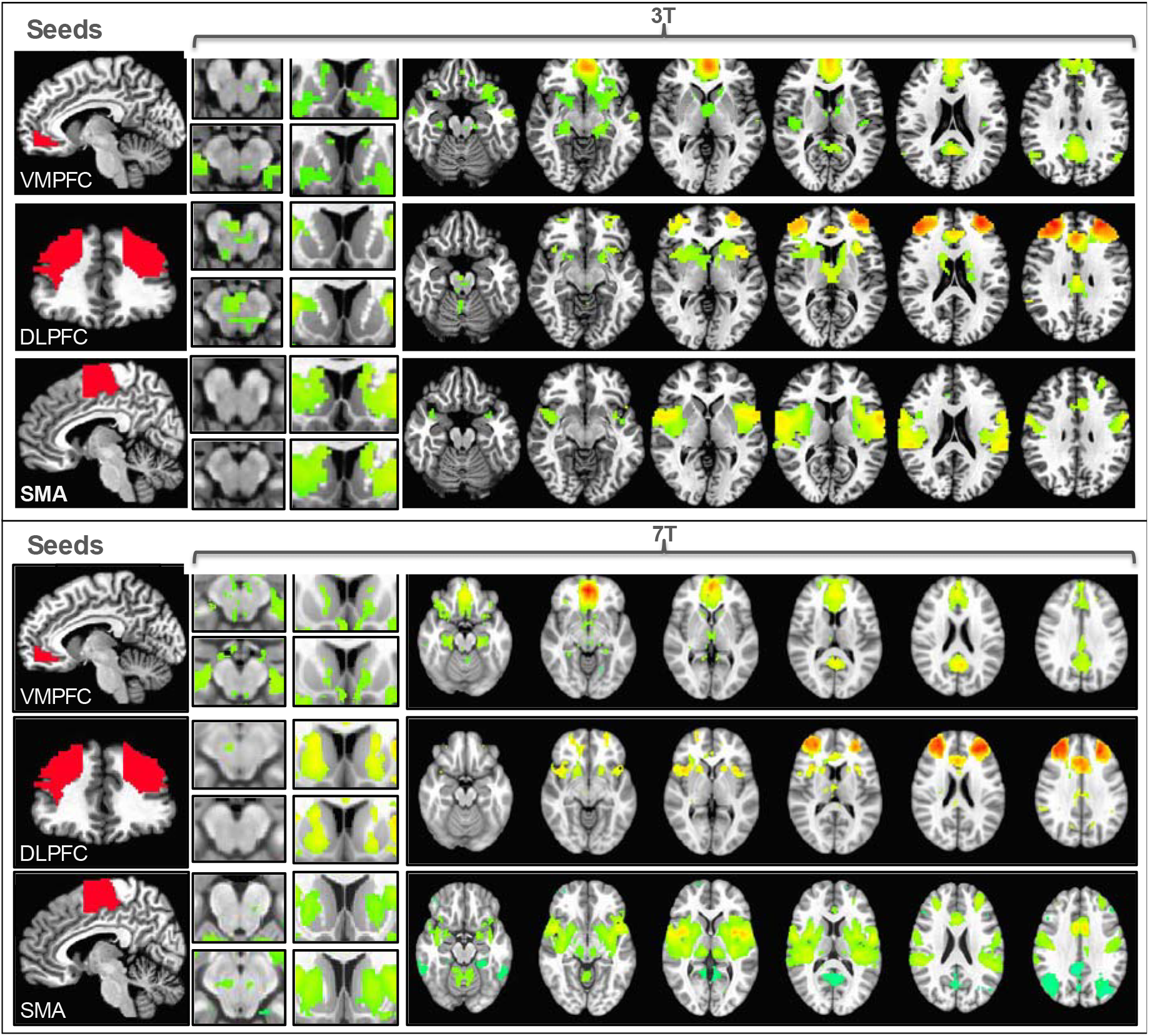
Functional connectivity of cortical-striatal-midbrain circuits with 3-Tesla (3T) and 7-Tesla (7T) functional magnetic resonance imaging (MRI). Resting state functional connectivity of three cortical seeds of interest (left, red) including ventromedial prefrontal cortex (VMPFC), dorsolateral prefrontal cortex (DLPFC), and supplementary motor area (SMA) was computed in healthy controls at 3T (top) and 7T (bottom). Connectivity maps are thresholded at p<0.0001 voxelwise for illustration purposes.

**Table 2.** Statistics of connectivity of seed regions of interest with 3-Tesla (3T) and 7-Tesla (7T) functional magnetic resonance imaging. Abbreviations: Ventromedial prefrontal cortex (VMPFC), dorsolateral prefrontal cortex (DLPFC), supplementary motor area (SMA), ventral tegmental area (VTA), posterior cingulate cortex (PCC), orbitfrontal cortex (OFC), anterior cingulate cortex (ACC), medial prefrontal cortex (mPFC), Z-score statistic (Z), whole-brain cluster corrected significance (corr), not significant (ns), cluster size (K), peak voxel coordinates (xyz).

| 3T |  |  |  |  |  |  | 7T |  |  |  |  |  |  |
| --- | --- | --- | --- | --- | --- | --- | --- | --- | --- | --- | --- | --- | --- |
| Region | Z | corr | K | x | y | z | Region | Z | corr | K | x | y | z |
| **VMPFC seed** |  |  |  |  |  |  | **VMPFC seed** |  |  |  |  |  |  |
| VMPFC | 14.3 | <0.05 | 7300 | 1.5 | -50.5 | -6 | VMPFC | 14.3 | <0.05 | 146283 | 1 | -52 | -5 |
| PCC | 7.2 | <0.05 | 1241 | -1.5 | 54.5 | 18 | PCC | 8.1 | <0.05 | 34842 | 4 | 56 | 19 |
| Cerebellum | 5.6 | <0.05 | 202 | -34.5 | 78.5 | -39 | OFC | 6.1 | <0.05 | 4089 | 37 | -31 | -15 |
| Lateral Parietal | 5.4 | <0.05 | 193 | 43.5 | 72.5 | 39 | Cerebellum | 5.4 | <0.05 | 3647 | 32 | 78 | -42 |
| Cerebellum | 7.3 | <0.05 | 169 | 31.5 | 75.5 | -36 | Lateral Parietal | -5.6 | <0.05 | 3593 | 29 | 53 | 68 |
| Lateral Parietal | 6.3 | <0.05 | 165 | -49.5 | 60.5 | 27 | Occipital | -5.7 | <0.05 | 2465 | 10 | 79 | 49 |
| Cerebellum | 5.9 | <0.05 | 152 | -7.5 | 54.5 | -45 | Occipital | -6.6 | <0.05 | 1883 | -40 | 81 | 18 |
| Cerebellum | 4.4 | <0.05 | 41 | -1.5 | 57.5 | -21 | Medial Parietal | -5.3 | <0.05 | 1475 | -10 | 68 | 60 |
| Occipital | 4.3 | ns | 21 | 25.5 | 99.5 | -12 | Lateral Parietal | -5.3 | <0.05 | 982 | -30 | 50 | 44 |
| Temporal | 4.1 | ns | 17 | 61.5 | 30.5 | -9 | Lateral Parietal | -4.9 | <0.05 | 946 | 55 | 37 | 43 |
| **DLPFC seed** |  |  |  |  |  |  | **DLPFC seed** |  |  |  |  |  |  |
| DLPFC | 10.9 | <0.05 | 9145 | -28.5 | -47.5 | 33 | DLPFC | 12.1 | <0.05 | 167874 | -31 | -50 | 26 |
| Lateral Parietal | 6.9 | <0.05 | 278 | -55.5 | 45.5 | 48 | Cerebellum | 8.9 | <0.05 | 9252 | 34 | 54 | -36 |
| Lateral Parietal | 6.5 | <0.05 | 155 | 52.5 | 54.5 | 48 | Cerebellum | 6.1 | <0.05 | 5887 | -40 | 52 | -37 |
| Cerebellum | 4.7 | <0.05 | 74 | -19.5 | 42.5 | -45 | Lateral Parietal | 5.8 | <0.05 | 4369 | 58 | 48 | 46 |
| Cerebellum | 4.2 | <0.05 | 36 | 10.5 | 45.5 | -42 | Lateral Parietal | 5.4 | <0.05 | 4284 | -57 | 37 | 46 |
| Medial Parietal | 5.4 | ns | 24 | -10.5 | 69.5 | 45 | Medial OFC | -4.8 | <0.05 | 1170 | 0 | -33 | -22 |
| Cerebellum | 4.1 | ns | 22 | -13.5 | 78.5 | -27 | Medial Parietal | 9.5 | <0.05 | 958 | -16 | 66 | 38 |
| Occipital | -4.3 | ns | 14 | -28.5 | 69.5 | -9 | Occipital | -4.9 | <0.05 | 942 | -16 | 92 | -2 |
|  |  |  |  |  |  |  | Medial Parietal | 4.1 | ns | 816 | 12 | 62 | 51 |
|  |  |  |  |  |  |  | Brainstem | 5.5 | ns | 674 | 7 | 44 | -60 |
| **SMA seed** |  |  |  |  |  |  | **SMA seed** |  |  |  |  |  |  |
| SMA | 11.9 | <0.05 | 9817 | -1.5 | 9.5 | 66 | SMA | 16.3 | <0.05 | 277708 | -1 | 8 | 63 |
| Lateral Parietal | -5.3 | <0.05 | 140 | -37.5 | 72.5 | -48 | Lateral Parietal | -13.4 | <0.05 | 17095 | -38 | 71 | 44 |
| Cerebellum | -6.8 | <0.05 | 116 | 43.5 | 66.5 | 42 | PCC | -7.1 | <0.05 | 13951 | -8 | 46 | 35 |
| DLPFC | 6.7 | <0.05 | 79 | 31.5 | -41.5 | 30 | Lateral Parietal | -10.6 | <0.05 | 11912 | 33 | 77 | 39 |
| Cerebellum | -4.7 | <0.05 | 64 | 37.5 | 69.5 | -48 | Cerebellum | 5.8 | <0.05 | 7978 | 16 | 45 | -25 |
| Cerebellum | 5.2 | <0.05 | 43 | 25.5 | 54.5 | -54 | DLPFC | -5.5 | <0.05 | 7850 | -31 | -9 | 45 |
| Lateral Parietal | -4.7 | <0.05 | 42 | -43.5 | 69.5 | 42 | DLPFC | 7.5 | <0.05 | 4739 | 32 | -45 | 23 |
| Motor Cortex | -6.2 | <0.05 | 40 | 40.5 | -17.5 | 48 | Thalamus | 14.3 | <0.05 | 4625 | -10 | 22 | 2 |
| PCC | -4.1 | <0.05 | 32 | -1.5 | 51.5 | 36 | Thalamus | 10.9 | <0.05 | 3803 | 8 | 24 | 2 |
| Thalamus | 4.3 | <0.05 | 27 | -10.5 | 18.5 | 0 | DLPFC | -5.9 | <0.05 | 2648 | 25 | -22 | 50 |
| **VTA seed** |  |  |  |  |  |  | **VTA seed** |  |  |  |  |  |  |
| VTA | 3.3 | <0.05 | 59 | -4.5 | 21.5 | -15 | VTA | 11.1 | <0.05 | 144215 | -2 | 19 | -16 |
|  |  |  |  |  |  |  | ACC/mPFC | 8.2 | <0.05 | 37531 | -4 | -27 | 16 |
|  |  |  |  |  |  |  | DLPFC | 4.9 | <0.05 | 1568 | 27 | -48 | 35 |
|  |  |  |  |  |  |  | VLPFC | 5.9 | ns | 528 | 20 | -53 | 6 |
|  |  |  |  |  |  |  | Cerebellum | 4.1 | ns | 505 | -33 | 56 | -33 |
|  |  |  |  |  |  |  | DLPFC | 4.3 | ns | 378 | -27 | -30 | 40 |
|  |  |  |  |  |  |  | Anterior Insula | 4.2 | ns | 264 | -41 | -9 | 9 |
|  |  |  |  |  |  |  | Medial Parietal | 4.2 | ns | 224 | -14 | 1 | 51 |
|  |  |  |  |  |  |  | Posterior Insula | 4.2 | ns | 197 | 45 | 21 | 10 |
|  |  |  |  |  |  |  | Medial Parietal | 4.6 | ns | 186 | 12 | 61 | 40 |

### Experiment 2: Reward-related neural network properties at high field in healthy controls and MDD

Subjects meeting DSM-5 criteria for MDD (N=15) were compared to the same HC cohort scanned at 3T described above (see Table 1 for demographics). A second cohort of patients meeting DSM-5 criteria for MDD (N=10) was compared to the same HC scanned at 7T described above (see Table 1 for demographics). Five of these MDD subjects completed both 7T and 3T scans. MDD patients had significantly higher depressive symptoms as measured by the Montgomery–Åsberg Depression Rating Scale (MADRS) and anhedonia as measured by the Snaith-Hamilton Pleasure Scale (SHAPS), compared to HC (p’s<0.001, see Table 1).

#### High field reveals latent neural network disturbances in depression

Firstly, VTA seed to voxel connectivity maps were compared at 3T and 7T in HC only. While at 3T, there was one significant cluster of VTA connectivity with neighbouring midbrain regions, higher field 7T provided more robust and detailed connectivity maps of the VTA, with medial PFC, ACC, DLPFC at the same whole-brain cluster-corrected thresholds (Fig 4A-B, Table 1 for statistics).

Given the specific interest in VTA connectivity with NAc and ACC (both subgenual and dorsal divisions) in MDD(18–20), we examined ROI-to-ROI connectivity between these regions. At 3T, there were no significant group differences between VTA-NAc or VTA-ACC (subgenual and dorsal) (p’s>0.05). However, at 7T, MDD subjects showed increased connectivity between VTA-subgenual ACC (p=0.009, Figure 4C). VTA-subgenual ACC connectivity in the MDD group positively correlated with the duration of the current episode (R=0.70, p=0.035) and the depressive symptom of anhedonia (R=0.67, p=0.036, Figure 4C). VTA-subgenual ACC connectivity was not correlated with age (p>0.05) or different based on gender (p>0.05) across the sample. There were no group differences in VTA-NAc connectivity or correlations between VTA-NAc connectivity and disorder severity or duration. Importantly, whole brain TSNR did not differ as a function of group between field strengths (Supp Figure 2).

**Figure 4.**
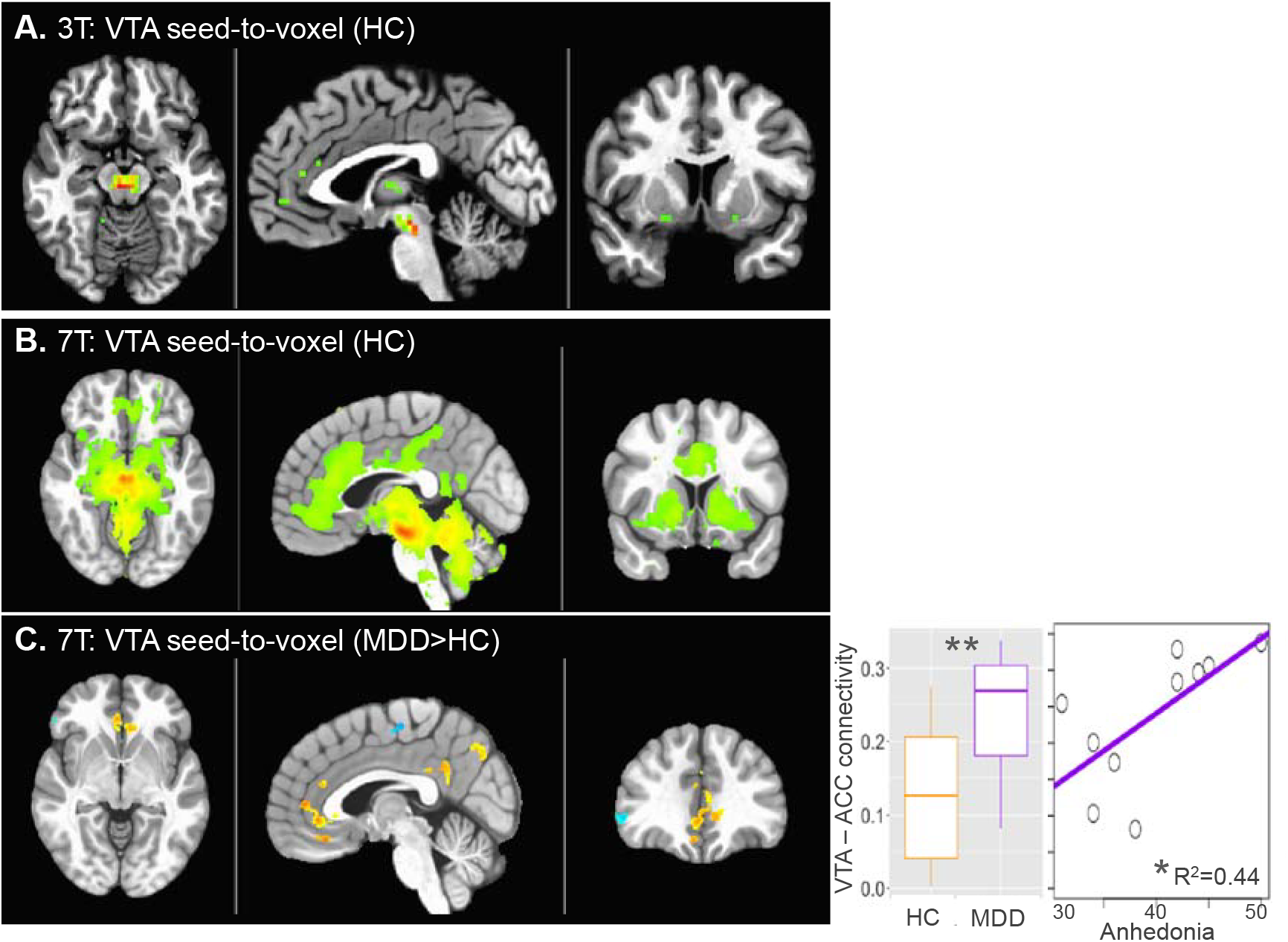
Functional connectivity of the ventral tegmental area (VTA) with 3-Tesla (3T) and 7-Tesla (7T) functional magnetic resonance imaging (MRI). Connectivity of the VTA with whole brain is shown for 3T (A) and 7T (B) in healthy controls (HC) (voxelwise p<0.001 for illustration). C. Patients with major depressive disorder (MDD) had higher connectivity of VTA with the anterior cingulate cortex (ACC) (p<0.01 voxelwise (Cluster>200) for illustration). VTA-ACC connectivity is plotted for MDD and HC and against anhedonia in the MDD group. **p<0.01; *p<0.05.

Whole-brain functional connectivity of the VTA was then compared between MDD patients and HC in an exploratory manner. At 3T, the comparison between MDD subjects (N=15) and HC (N=17 from Experiment 1) revealed no clusters at voxelwise p<0.01, Cluster>100. At 7T, the comparison between a smaller sample of MDD subjects (N=10) and HC (N=17 from Experiment 1) revealed several clusters at voxelwise p<0.01, Cluster>100, with increased connectivity with ACC in MDD as the top cluster (Fig 4C, Table 2). While this observation corroborates the ROI-to-ROI based analysis, it did not reach whole brain cluster-corrected significance in this sample (Cluster>4645 for alpha<0.05) and should therefore be interpreted with some caution.

## Discussion

We report substantially improved signal power across the brain with ultra-high field 7T fMRI compared to 3T that is boosted by multi-echo based denoising. We observed an increase in whole-brain TSNR of approximately 230% at 7T compared to 3T. Functional connectivity between diverse brain regions relevant to MDD and other neuropsychiatric disorders was significantly enhanced at 7T compared to 3T, including of VTA, a small midbrain structure that is limited by poor TSNR at lower clinical field strength. Furthermore, hyperconnectivity between the VTA and subgenual ACC in MDD was revealed at 7T, consistent with pre-clinical findings in a validated rodent model of depression(19, 20), while this alteration was not observed at 3T. Together, this work indicates the considerable utility of ultra-high field 7T for characterizing pathological alterations in neural architecture relevant to neuropsychiatric disorders.

While clinical applications of ultra-high field 7T neuroimaging have been posited for multiple disorders including stroke, epilepsy, multiple sclerosis(6), there has been less emphasis on more subtle psychiatric disorders such as MDD. This work adds to other demonstrations of the utility of ultra-high field functional MRI, including comparisons of task based functional MRI (38) and of the resting state connectivity of the Habenula (39). Determining biomarkers for depression and other psychiatric disorders will be crucial for the successful advancement of diagnostics and targeted, precision medicine. Ultra-high field MRI may provide additional pathophysiological insight for early disorder detection and early intervention.

The VTA dopaminergic projections are the most well-established characterization of the ‘reward circuit’(18). While there are no other 7T fMRI studies of VTA in humans with MDD, the current demonstration of VTA hyperconnectivity in MDD is in line with several pre-clinical models of depression. A well-validated rodent model of chronic social defeat stress is characterized by reduced social interaction and reduced sucrose preference, indicating an anhedonic phenotype in susceptible animals (18–20). This model is also characterized by VTA hyperactivity, and normalizing VTA hyperconnectivity reduces the depressive, anhedonic phenotype(40). Another recent preclinical study has demonstrated that VTA connectivity with ACC mediates motivation for rewards(41) and heightened subgenual ACC connectivity has been previously demonstrated in patients with MDD(42). The current study confirms the observation of VTA hyperconnectivity in humans with depression who are free of medication. Future studies should examine the functional organization of the broader cortical-subcortical reward circuit as a whole, which encompasses other neural substrates in the PFC, hippocampus and amygdala, as well as their roles in discerning and responding to environmental signals of reward, punishment and threat, in a larger sample size.

Issues around patient contraindication, dizziness and claustrophobia are heightened at 7T, meaning not all patients will be eligible. There also remain issues in B0 and B1 inhomogeneity, resulting in distortion and drop-out, respectively, requiring advanced shimming and specialized pulse sequence designs(6). The current study has several limitations. One specific limitation is that the voxel size was different between 7T (2.5mm^3^) compared to 3T (3mm^3^). Voxel size contributes to connectivity estimates, particularly for small nuclei such as the VTA (43). Other ultra-high field MRI protocols have used smaller voxel sizes, which related to improved connectivity estimates in motor regions although this may not be the case for other regions (43). The precision of the whole-brain functional connectivity maps demonstrated may be influenced by the slightly higher resolution (smaller voxel size) of the 7T protocol compared to 3T. The current 7T voxel size is also relatively large compared to other studies (38, 39), a trade-off required for the use of the multi-echo functional MRI acquisition protocol. The TR was also longer at 7T compared to 3T, meaning the number of frames was higher for 3T for the same scan time. This is expected to enhance the temporal resolution of the 3T scans compared to 7T. Finally, the iPAT acceleration factor was higher at 7T compared to 3T, which would be expected to influence SNR. Therefore while we demonstrate improvements in multiple different measurements, including TSNR, connectivity metrics and whole-brain connectivity maps for both the same subjects and different subjects scanned with 3T and 7T, some findings may be influenced by differences in acquisition protocols.

We provide a characterization of the utility of ultra-high field 7-Tesla MRI for applications in functional connectivity mapping of neural structures that are critical to the pathophysiology of neuropsychiatric disorders. We emphasize the use of multi-echo based acquisition and denoising methods at ultra-high field.

## Funding and Disclosures

Funding was provided by R21MH109771 and R01MH109544. Additional support was provided by the Friedman Brain Institute and by the Ehrenkranz Laboratory for Human Resilience, both components of the Icahn School of Medicine at Mount Sinai. The authors disclose no conflicts of interest.

## References

1. Veer IM, Beckmann CF, van Tol MJ, Ferrarini L, Milles J, Veltman DJ, et al. (2010): Whole brain resting-state analysis reveals decreased functional connectivity in major depression. Front Syst Neurosci. 4.

2. Sheline YI, Price JL, Yan Z, Mintun MA (2010): Resting-state functional MRI in depression unmasks increased connectivity between networks via the dorsal nexus. Proc Natl Acad Sci U S A. 107:11020–11025.

3. Zeng LL, Shen H, Liu L, Wang L, Li B, Fang P, et al. (2012): Identifying major depression using whole-brain functional connectivity: a multivariate pattern analysis. Brain. 135:1498–1507.

4. Anand A, Li Y, Wang Y, Wu J, Gao S, Bukhari L, et al. (2005): Activity and connectivity of brain mood regulating circuit in depression: a functional magnetic resonance study. Biol Psychiatry. 57:1079–1088.

5. Beisteiner R, Robinson S, Wurnig M, Hilbert M, Merksa K, Rath J, et al. (2011): Clinical fMRI: evidence for a 7T benefit over 3T. Neuroimage. 57:1015–1021.

6. Balchandani P, Naidich TP (2015): Ultra-High-Field MR Neuroimaging. Am J Neuroradiol. 36:1204–1215.

7. Triantafyllou C, Hoge RD, Krueger G, Wiggins CJ, Potthast A, Wiggins GC, et al. (2005): Comparison of physiological noise at 1.5 T, 3 T and 7 T and optimization of fMRI acquisition parameters. Neuroimage. 26:243–250.

8. van der Zwaag W, Francis S, Head K, Peters A, Gowland P, Morris P, et al. (2009): fMRI at 1.5, 3 and 7 T: characterising BOLD signal changes. Neuroimage. 47:1425–1434.

9. Sladky R, Baldinger P, Kranz GS, Trostl J, Hoflich A, Lanzenberger R, et al. (2013): High-resolution functional MRI of the human amygdala at 7 T. Eur J Radiol. 82:728–733.

10. Schenck JF (1996): The role of magnetic susceptibility in magnetic resonance imaging: MRI magnetic compatibility of the first and second kinds. Medical physics. 23:815–850.

11. Windischberger C, Langenberger H, Sycha T, Tschernko EA, Fuchsjager-Mayerl G, Schmetterer L, et al. (2002): On the origin of respiratory artifacts in BOLD-EPI of the human brain. Magn Reson Imaging. 20:575–582.

12. Merboldt KD, Fransson P, Bruhn H, Frahm J (2001): Functional MRI of the human amygdala? NeuroImage. 14:253–257.

13. Robinson S, Windischberger C, Rauscher A, Moser E (2004): Optimized 3 T EPI of the amygdalae. NeuroImage. 22:203–210.

14. Collins PY, Patel V, Joestl SS, March D, Insel TR, Daar AS, et al. (2011): Grand challenges in global mental health. Nature. 475:27–30.

15. Keedwell PA, Andrew C, Williams SC, Brammer MJ, Phillips ML (2005): The neural correlates of anhedonia in major depressive disorder. Biol Psychiatry. 58:843–853.

16. Pizzagalli DA, Holmes AJ, Dillon DG, Goetz EL, Birk JL, Bogdan R, et al. (2009): Reduced Caudate and Nucleus Accumbens Response to Rewards in Unmedicated Individuals With Major Depressive Disorder. Am J Psychiat. 166:702–710.

17. Wohlschlaeger A, Karne, H., Jordan, D., Lowe, M., Jones, S. E., & Anand, A. (2018): Spectral Dynamics of Resting State fMRI within the Ventral Tegmental Area and Dorsal Raphe Nuclei in Medication-free Major Depressive Disorder in Young Adults. Frontiers in Psychiatry. 9:163.

18. Russo SJ, Nestler EJ (2013): The brain reward circuitry in mood disorders. Nature Reviews Neuroscience. 14:609–625.

19. Chaudhury D, Walsh JJ, Friedman AK, Juarez B, Ku SM, Koo JW, et al. (2013): Rapid regulation of depression-related behaviours by control of midbrain dopamine neurons. Nature. 493:532-+.

20. Friedman AK, Walsh JJ, Juarez B, Ku SM, Chaudhury D, Wang J, et al. (2014): Enhancing Depression Mechanisms in Midbrain Dopamine Neurons Achieves Homeostatic Resilience. Science. 344:313–319.

21. Groenewegen HJ, Galis-de Graaf Y, Smeets WJAJ (1999): Integration and segregation of limbic cortico-striatal loops at the thalamic level: an experimental tracing study in rats. J Chem Neuroanat. 16:167–185.

22. Groenewegen HJ, Wright CI, Beijer AVJ, Voorn P (1999): Convergence and segregation of ventral striatal inputs and outputs. Ann Ny Acad Sci. 877:49–63.

23. Haber SN (2003): The primate basal ganglia: parallel and integrative networks. J Chem Neuroanat. 26:317–330.

24. First MW, JBW; Karg, RS; Spitzer, RL Structured Clinical Interview for DSM-5—Research Version (SCID-5 for DSM-5, Research Version; SCID-5-RV). Arlington, VA, American Psychiatric Association, 2015.

25. Montgomery SA ÅM (1979): A new depression scale designed to be sensitive to change. Br J Psychiatry. 134:382–389.

26. Snaith RP, Hamilton M, Morley S, Humayan A, Hargreaves D, Trigwell P (1995): A scale for the assessment of hedonic tone the Snaith-Hamilton Pleasure Scale. Br J Psychiatry. 167:99–103.

27. Marques JP, Kober T, Krueger G, van der Zwaag W, Van de Moortele PF, Gruetter R (2010): MP2RAGE, a self bias-field corrected sequence for improved segmentation and T1-mapping at high field. Neuroimage. 49:1271–1281.

28. Marques JP, Gruetter R (2013): New developments and applications of the MP2RAGE sequence--focusing the contrast and high spatial resolution R1 mapping. Plos One. 8:e69294.

29. Kundu P, Inati SJ, Evans JW, Luh WM, Bandettini PA (2012): Differentiating BOLD and non-BOLD signals in fMRI time series using multi-echo EPI. Neuroimage. 60:1759–1770.

30. Kundu P, Voon V, Balchandani P, Lombardo MV, Poser BA, Bandettini PA (2017): Multi-echo fMRI: A review of applications in fMRI denoising and analysis of BOLD signals. Neuroimage. 154:59–80.

31. Avants BB, Epstein CL, Grossman M, Gee JC (2008): Symmetric diffeomorphic image registration with cross-correlation: evaluating automated labeling of elderly and neurodegenerative brain. Med Image Anal. 12:26–41.

32. Avants BB, Tustison NJ, Song G, Cook PA, Klein A, Gee JC (2011): A reproducible evaluation of ANTs similarity metric performance in brain image registration. Neuroimage. 54:2033–2044.

33. Cox RW (1996): AFNI: software for analysis and visualization of functional magnetic resonance neuroimages. Comput Biomed Res. 29:162–173.

34. Morris LS, Kundu P, Dowell N, Mechelmans DJ, Favre P, Irvine MA, et al. (2016): Fronto-striatal organization: Defining functional and microstructural substrates of behavioural flexibility. Cortex. 74:118–133.

35. Taylor PA, Saad ZS (2013): FATCAT: (an efficient) Functional and Tractographic Connectivity Analysis Toolbox. Brain Connect. 3:523–535.

36. Cox RW, Chen G, Glen DR, Reynolds RC, Taylor PA (2017): fMRI clustering and false-positive rates. Proc Natl Acad Sci U S A. 114:E3370–E3371.

37. Patel AX, Bullmore ET (2016): A wavelet-based estimator of the degrees of freedom in denoised fMRI time series for probabilistic testing of functional connectivity and brain graphs. Neuroimage. 142:14–26.

38. Torrisi S, Chen G, Glen D, Bandettini PA, Baker CI, Reynolds R, et al. (2018): Statistical power comparisons at 3T and 7T with a GO / NOGO task. Neuroimage. 175:100–110.

39. Torrisi S, Nord CL, Balderston NL, Roiser JP, Grillon C, Ernst M (2017): Resting state connectivity of the human habenula at ultra-high field. Neuroimage. 147:872–879.

40. Friedman AK, Juarez B, Ku SM, Zhang H, Calizo RC, Walsh JJ, et al. (2016): KCNQ channel openers reverse depressive symptoms via an active resilience mechanism. Nat Commun. 7:11671.

41. Elston TW, Bilkey DK (2017): Anterior Cingulate Cortex Modulation of the Ventral Tegmental Area in an Effort Task. Cell Rep. 19:2220–2230.

42. Greicius MD, Flores BH, Menon V, Glover GH, Solvason HB, Kenna H, et al. (2007): Resting-state functional connectivity in major depression: abnormally increased contributions from subgenual cingulate cortex and thalamus. Biol Psychiatry. 62:429–437.

43. Newton AT, Rogers BP, Gore JC, Morgan VL (2012): Improving measurement of functional connectivity through decreasing partial volume effects at 7 T. Neuroimage. 59:2511–2517.

